# Targeted sex ratio distortion in the mouse

**DOI:** 10.1101/2025.06.19.660540

**Authors:** Hermann Bauer, Frederic Koch, Bettina Lipkowitz, Jürgen Willert, Gaby Bläß, Manuela Scholze-Wittler, Sandra Währisch, Lars Wittler, Bernhard G. Herrmann

## Abstract

In animal breeding, female or male progeny is preferred depending on the use of the animals. Especially in livestock, offspring with undesired sex is of low economic value and often culled. Here we achieve targeted sex ratio distortion in the mouse, leveraging components of a selfish supergene, the *t*-haplotype. Utilizing its central element, the *t*-responder, we generate a highly “selfless” genetic element disabling transgenic sperm. When integrated on the X or Y chromosome it results in a high prevalence of non-transgenic offspring of the desired sex.

Our strategy overcomes key limitations of existing strategies, including compromised animal health, reduced fertility, and fully transgenic breeding stocks, which might foster acceptance of its application to livestock. The efficacy of our approach marks a groundbreaking advancement in animal breeding.

**Graphical abstract:** 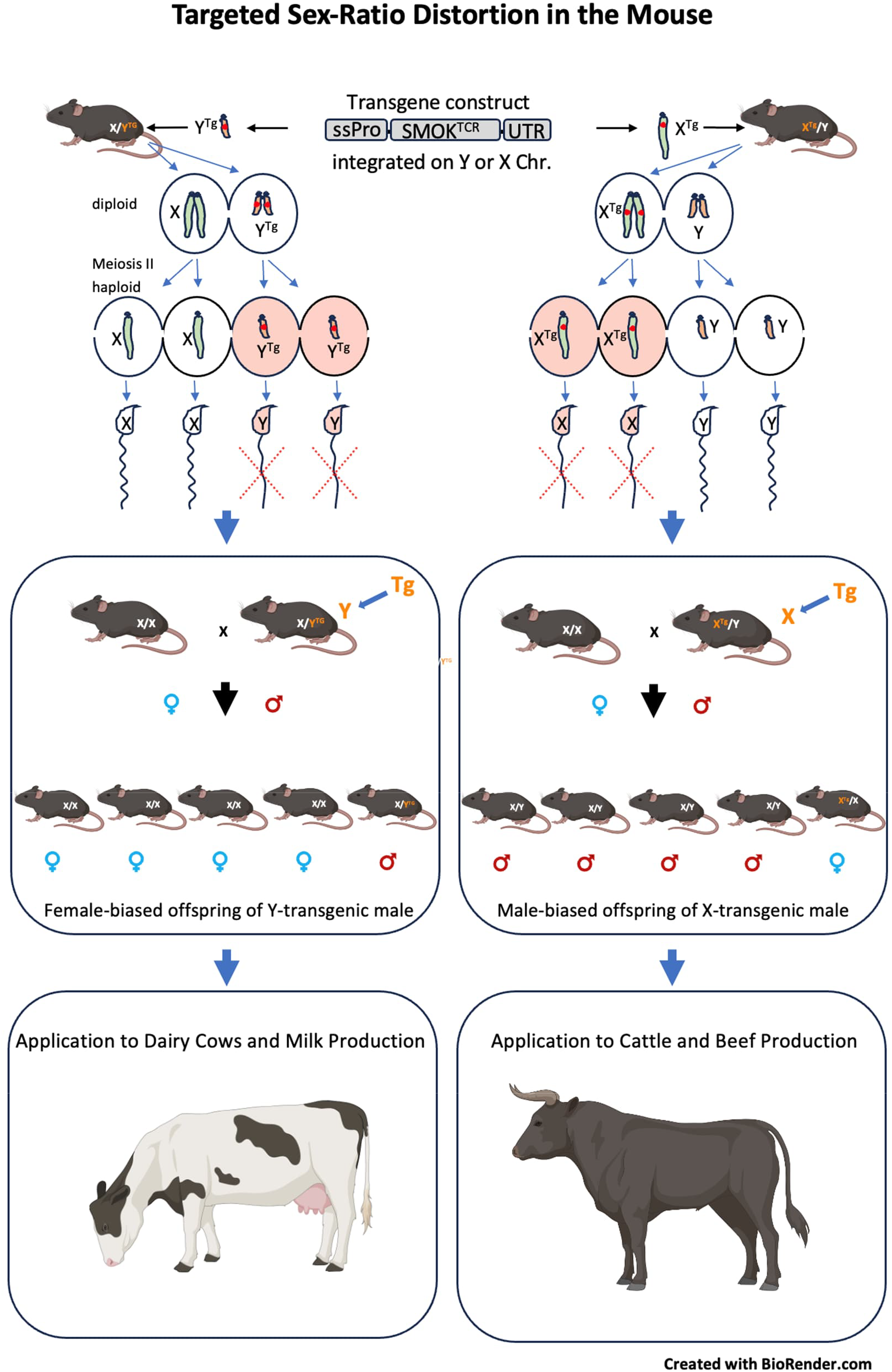

**Transgene integration on X or Y chromosome affects the motility of transgenic sperm resulting in a strong sex-bias towards non-transgenic/wildtype offspring. Application to livestock can reduce unwanted progeny and transform farm animal breeding.**

## Introduction

Animal production faces major challenges by limited resources for a growing human population and increasing standards for animal welfare. A substantial improvement in breeding efficiency and mitigation of ethical problems could be achieved by preferential inheritance of desired genetic traits. This is particularly relevant in livestock breeding, where almost half of the progeny with undesired sex often has little or no economic value and is culled, leading to both economic and ethical issues (Douglas and Turner 2020). For instance, female progeny is preferred in dairy cattle, with only a few males needed as semen producers. Conversely, in beef cattle, male offspring is valued for meat production, while females are primarily used for breeding.

Controlling the sex of progeny therefore is of paramount interest. Several approaches for altering the sex ratio have been developed with pioneering work conducted in the mouse model. These include transgenic sex reversal by Sry knock-out or sex chromosome shredding (Zuo, Huo et al. 2017), often resulting in animals with reproductive defects (Zuo, Huo et al. 2017) (Kurtz, Lucas-Hahn et al. 2021). Attempts to bias the sex ratio of offspring in the mouse by X-chromosome shredding in spermatogenic cells have failed recently (Bunting, Godahewa et al. 2024).

Gene drives, while theoretically applicable for inducing transgenic sex reversal, have shown low efficiency in mammals, highlighting the challenges of adapting strategies successfully used for insects to mammals (Grunwald, Gantz et al. 2019) (Weitzel, Grunwald et al. 2021). Other methods for shifting the sex ratio involve incubation of sperm with antibodies against a Y-sperm specific epitope, sperm pre-treatment with undisclosed substances (reviewed in (Quelhas, Pinto-Pinho et al. 2023) or manipulation of Toll-like receptors in vitro (Umehara, Tsujita et al. 2019, Ren, Xi et al. 2021). These methods require extensive manipulation of sperm samples and to some degree are controversial with respect to their effectiveness. In cattle, fluorescence-activated cell sorting (FACS) is used to separate X-and Y-chromosome-bearing sperm for artificial insemination (Holden and Butler 2018) (Obuchi, Osada et al. 2019). However, sperm sorting technology is costly for the breeder and not well established in other livestock species. In a recent approach in the mouse, selective lethality of conceptuses carrying the Y chromosome is induced by the combined inheritance of an autosomal Cas9 transgene and a Y-chromosomal guide-RNA cassette directed against essential embryonic genes (Yosef, Edry-Botzer et al. 2019). Due to incomplete penetrance, some animals develop to term and are malformed, raising animal welfare concerns. Another more recently developed synthetic lethal system induces embryonic death of all progeny with undesired sex prior to implantation (Douglas, Maciulyte et al. 2021). However, all parents and offspring are transgenic and the productivity of the breeding stock is strongly reduced.

A genetic system designed towards generating a surplus of non-transgenic progeny has been presented recently in a pre-print (Yosef, Mahata et al. 2023). It employs dCas9 mediated suppression of a gene required for spermatid maturation in mouse impairing the fertilization capability of transgenic sperm. However, transgenic males are growth impaired and show strongly reduced fertility in natural breeding, though their sperm is effective in in vitro fertilization.

Several naturally occurring genetic systems influencing transmission ratio in their favor have been reported (Burt and Trivers 2006), some of which hold promise for a practical application. The most prominent element causing transmission ratio distortion in mammals is the mouse *t*-haplotype, a naturally occurring variant of chromosome 17, able to strongly increase its transmission from t/+ heterozygous males to their offspring at the expense of the wild type chromosome (Lyon 2003). The *t*-haplotype achieves high transmission by a “killer-antidote” mechanism. Its “killer” genes, so-called *t*-distorters, act harmfully by affecting progressive movement in all sperm (Bauer, Willert et al. 2005, Bauer, Veron et al. 2007, Bauer, Schindler et al. 2012, Amaral and Herrmann 2021). The “antidote”, termed *t*-complex-responder (Tcr), encodes a dominant-negative form of Smok (Smok-Tcr), able to rescue progressive motility. But it does so only in sperm carrying the gene providing them with a fertilization advantage over their competitors (Herrmann, Koschorz et al. 1999).

However, in the absence of *t*-distorters, the dominant negative action of Smok-Tcr results in a disadvantage of sperm. This low transmission ratio can be phenocopied by Tcr transgenes.

We have been the first reporting successful sex ratio distortion in a mammal (Herrmann, Koschorz et al. 1999). Using a Tcr trangene randomly integrated on the Y chromosome we achieved a surplus of male offspring at a 2:1 (male/female) ratio, but only in combination with *t*-distorters. In contrast, in the present study we used single copy integrations of Tcr transgenes into defined landing sites on the X or Y chromosome. This allowed manipulation of the sex ratio by design. We optimized the Tcr transgene to further increase its efficiency from 65% to up to 88% non-transgenic offspring of the desired sex, again without using distorters. This is the first report demonstrating highly efficient sex ratio distortion in mice in favor of either male or female non-transgenic progeny.

## Results

### Achieving sex ratio distortion while maintaining health and fertility

To achieve targeted sex-ratio distortion without affecting the health or fertility of the animals several conditions have to be met. Transgene expression should be restricted to transgenic sperm and therefore occur post-meiotically in haploid spermatids only. To achieve that, the promoter driving transgene expression should be spermatid-specific, and transgene-derived mRNA and proteins should be retained in the transgenic spermatids, a challenging requirement given that spermatids typically share mRNA due to their development in a syncytium (Braun, Behringer et al. 1989). Notably, *Smok-Tcr* is the only case where non-sharing has been demonstrated experimentally (Veron, Bauer et al. 2009). In addition, transgene activity in somatic cells which might compromise the health of the animal, should be avoided. Moreover, the expression level of the transgene on a sex chromosome must be high enough to achieve an effect. Since the X and the Y chromosome undergo meiotic sex chromosome inactivation (MSCI) resulting in silencing of many regions (Turner 2007), transgene expression levels may be affected. Too high expression, on the other hand, might allow sharing of Tg-derived mRNA between spermatids to some degree, which might reduce the TRD effect. Furthermore, the transgenic construct should not interfere directly or via chromatin silencing with the expression of an endogenous gene which might compromise the health or fertility of the transgenic male. Hence, integration into transcription units should be avoided.

To achieve targeted inheritance of desired genetic traits we first established site directed, single copy and selection-marker free integration of *Smok-Tcr* constructs. Such transgenes when integrated in the vicinity of the *Smok2b* locus showed TRD to a similar degree as randomly integrated transgenes (data not shown). To achieve sex ratio distortion, we integrated *Smok-Tcr* transgenes controlled by autosomal promoters on the X and Y chromosome. However, these transgenes showed very low expression and no sex ratio distortion (data not shown). We reasoned that this might be due to silencing of transgene promoters by chromatin adjacent to the transgene integration sites.

### Finding landing sites on the X and Y chromosome escaping MSCI

Finding an appropriate landing site outside gene bodies and active during spermiogenesis is challenging. We looked for landing sites in regions that become active during spermiogenesis, based on epigenetic chromatin signatures. To pinpoint such regions, we performed Chromatin Immunoprecipitation Sequencing (ChIP-seq) experiments on nuclear material isolated from mouse testis at three time points after birth (post partum; p.p.): day (d) 12, 16, 24, and from adult testis. The first wave of spermatogenesis starts right after birth, and therefore this first differentiation process can be staged. Day12 marks the meiotic stage, day 16 the onset of spermiogenesis (the haploid stage), and day 24 a later stage of spermiogenesis in which a particular set of genes including members of the Smok gene family are first expressed. Additionally, we performed RNA-sequencing (RNA-seq) of staged testes (Table 1). The combination of ChIP-seq and RNA-seq data allowed us to identify suitable landing sites. We chose a region upstream of *Zfy2* on the Y chromosome (LS-*Zfy2*), which becomes active as indicated by H3K27 acetylation (upregulated between d24 and adult; Fig. 1a; Table 1).

**Table 1.** Temporal expression profile of post-meiotically activated genes determined by RNA-seq of staged testis samples. Numbers are FPKM values.

| **Gene** | **day 12 p.p.** | **day 16 p.p.** | **day 24 p.p.** | **adult** |
| --- | --- | --- | --- | --- |
| *Smok2a* | 0 | 0 | 1.11 | 6.49 |
| *Smok2b* | 0 | 0 | 1.04 | 5.62 |
| *Spermatid expressed gene* | 0 | 0.34 | 13.99 | 55.96 |
| *Akap4* | 0.27 | 0.11 | 97.65 | 567.03 |
| *Zfy2* | 5.28 | 5.26 | 4.80 | 9.88 |

**Figure 1.**
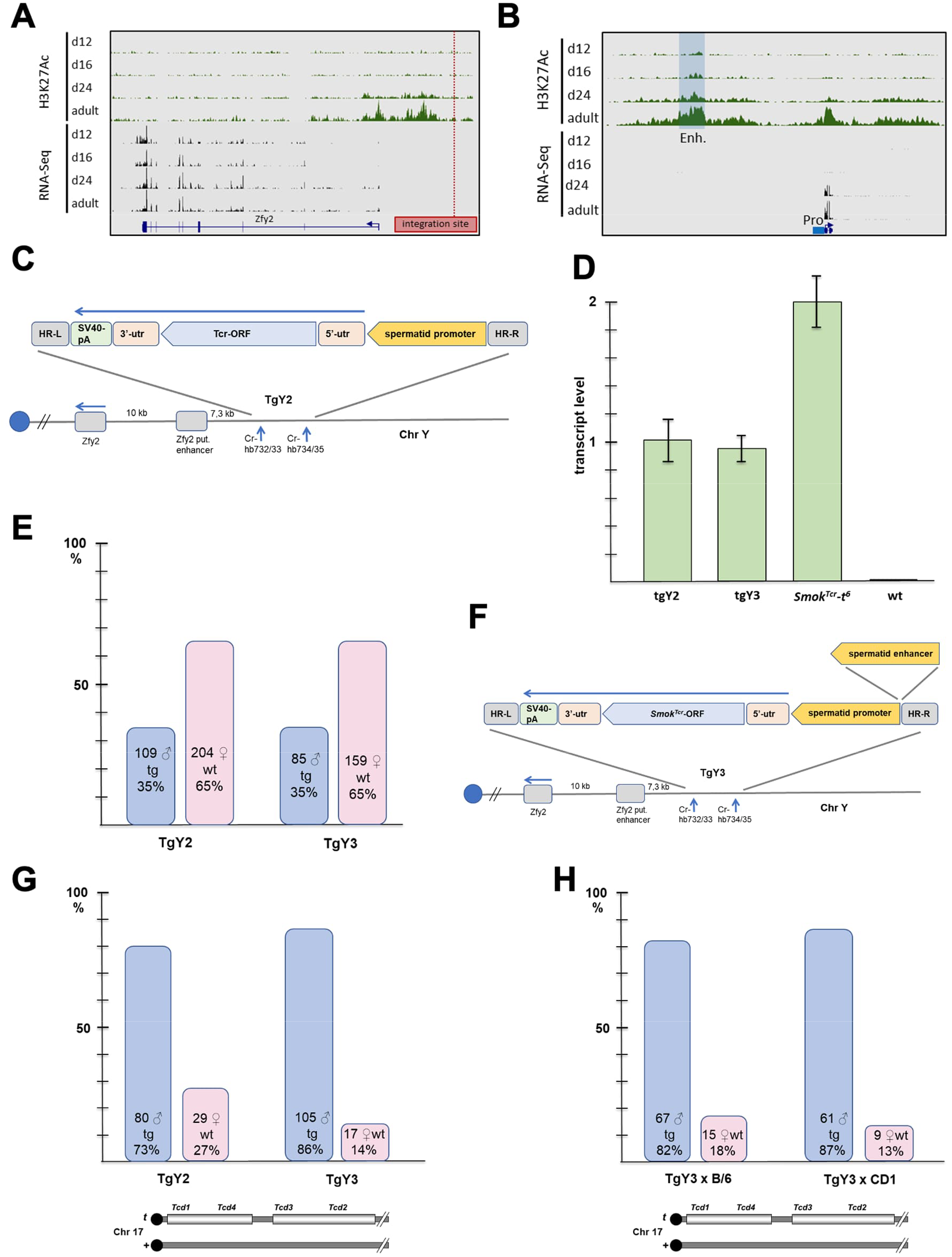
Targeted sex ratio distortion by a transgene integrated on the Y chromosome. (**A**) Identification of a suitable integration site near the *Zfy2* gene based on histone modification data. The H3K27ac signatures and expression profiles in post-partum testes indicate gene expression from an upstream exon from d24 on and identify nearby active chromatin regions and a landing site for our transgenes. (**B**) ChIP-seq and RNA-seq identify a spermatid promoter and spermatid enhancer for transgene expression. (**C**) TgY2, a *Smok-Tcr* transgene integrated into the landing site near *Zfy2* identified in (A). (**D**) Analysis of Tg expression by qPCR, in comparison to endogenous *Smok-Tcr*. (**E**) Tg transmission and sex ratio in offspring from males carrying the indicated Tg construct. (**F**) TgY3, a *Smok-Tcr* transgene carrying in addition a putative spermatid-specific enhancer integrated into the landing site near *Zfy2*. (**G**) Analysis of Tg transmission and sex ratio in offspring from males carrying *t*-distorters encoded on the *t*-haplotype *t*^*h51*^-*t*^*h18*^ on chromosome 17 in addition to the indicated Tg construct on the Y chromosome. (**H**) TgY3 transmission and sex ratio after backcrossing to different genetic backgrounds (B/6 or CD1).

### A post-meiotically active promoter and enhancer escaping MSCI suitable for post-meiotic transgene expression

We hypothesized that a promoter escaping X/Y inactivation and becoming active on the X or Y chromosome in late spermiogenesis should be better suited for achieving stage-specific transgene expression at a sufficiently high level than the autosomal *Smok-Tcr* promoter. To identify such a promoter, we searched for genes expressed at moderate levels starting from d24 p.p. and exceeding *Smok2a*/*Smok2b* expression levels. Since most Y expressed genes activated during spermiogenesis are repetitive and promoter strength is therefore difficult to assign to a particular gene locus, we focused on single copy genes located on the X chromosome.

We identified a promoter that fulfilled all criteria discussed above. It is spermiogenesis-specific and expressed from day 24 p.p. at a moderate level (Fig. 1b, Table 1). This promoter was used to drive the expression of a *Smok-Tcr* coding sequence flanked by a 5′- and 3′-UTR and the SV40 poly(A) signal (TgY2; Fig. 1c). Using CRISPR/Cas9, we integrated the transgene into LS-*Zfy2* in mouse ES cells, generated mice via morula aggregation, and evaluated stage- and site-specific transgene expression via qPCR of testis RNA.

The transgene displayed restricted spermatid-specific expression, albeit at approximately half the expression level of the *t*-haplotype *Smok-Tcr* (Fig. 1d). This nonetheless caused significant sex ratio distortion favoring non-transgenic females (65%; Fig. 1e). We aimed to increase expression of the transgene by adding an enhancer sequence. Using our ChIP-seq data we identified a putative enhancer region located upstream of the spermatid-expressed gene, which becomes active when the gene is first transcribed, as based on histone modification data (H3K27Ac+), indicating active enhancers (Fig. 1b). We integrated the putative enhancer sequence into TgY2 upstream of the promoter driving *Smok-Tcr* expression and generated the mouse line TgY3 (Fig. 1f). TgY3 males showed the same expression level as TgY2 males and generated 65% wildtype female offspring, just like TgY2 (Fig. 1d,e).

We crossed both lines to a sensitized genetic background containing a partial *t*-haplotype, *t*^*h51*^/*t*^*h18*^ expressing *t*-distorter genes causing high transmission of *Smok-Tcr* alleles (Lyon 1984). TgY3 showed significantly higher transmission than TgY2 (86% vs 73% males, Fisher exact test statistic value 0.0205, Fig. 1g, Table 2).

**Table 2.** Tg-transmission and sex-ratio distortion. Transmission rates of *Smok-Tcr*-transgenes (tg/0; +/+) without or in presence of distorter factors (tg/0; *t*^*h51*^/*t*^*h18*^). We performed a two-tailed binomial test, comparing to expected Mendelian chromosome transmission (50%). Abbr.: G0 stands for founder generation, N for backcross generation.

| **Line** | **tg** | **Non- tg** | **total** | **%tg** | **%non-tg females** | **Binomial test** |
| --- | --- | --- | --- | --- | --- | --- |
| tgY2; +/+ | 109 | 204 | 313 | 35 | 65 | 8,64939E-08 |
| tgY2/0; $t^{h51}/t^{h18}$ N1 | 80 | 29 | 109 | 73 | 27 | 1,08091E-06 |
| tgY3/0; +/+ G0 | 85 | 159 | 244 | 35 | 65 | 2,50483E-06 |
| tgY3/0; $t^{h51}/t^{h18}$ N1 | 105 | 17 | 122 | 86 | 14 | 1,14746E-16 |
| tgY3/0; $t^{h51}/t^{h18}$<br>N2, N3, N4 x B/6 | 67 | 15 | 82 | 82 | 18 | 5,25732E-09 |
| tgY3/0; $t^{h51}/t^{h18}$<br>N1, N2, N3 x CD1 | 61 | 9 | 70 | 87 | 13 | 1,28439E-10 |
| tgY4/0; G0 | 25 | 181 | 206 | 12 | 88 | 2,23574E-30 |
| **Line** | **tg** | **Non- tg** | **total** | **%tg** | **% non-tg males** | **Binomial test** |
| tgX2/0; G0 | 53 | 188 | 241 | 22 | 78 | 6,66766E-19 |
| **Controls** | **males** | **females** | **total** |  | **% males** |  |
| Offspring from control G0 males | 278 | 267 | 545 |  | 51 | 0,668433241 |
| Offspring from control N1, N2, N3 x B/6 males | 214 | 201 | 415 |  | 52 | 0,555875244 |

TRD of *t*-haplotypes has been shown to be highly variable depending on the genetic background (Gummere, McCormick et al. 1986). Therefore, we backcrossed TgY3 together with *t*^*h51*^/*t*^*h18*^ to the inbred strain C57BL/6J (B/6) and to an outbred strain with high fertility, CD1. Transmission tests yielded 82% and 87% male offspring, respectively, indicating similarly high effects of both transgenes on either background.

The combined data showed that both transgene constructs are effectively causing SRD, either in favor or at the disadvantage of the Tg-carrying chromosome. The effect is stable through generations and preserved on different genetic backgrounds (Table 2). The rescue effect of Smok-Tcr in the presence of *t*-distorters generated more pronounced deviations from the Mendelian sex ratio than *Smok-Tcr* transgenes in a wild-type background.

### Optimizing *Smok-Tcr* improves the sex bias in progeny

The most wanted practical application of SRD is meiotic drag of a transgene integrated on the unwanted sex chromosome, resulting in high transmission of the preferred, non-transgenic sex chromosome. However, as shown above, this “low-ratio” phenotype is harder to obtain than high transmission. Therefore, we explored if transgene transmission could be further lowered by increasing *Smok-Tcr* activity. To this end we optimized TgY3 to obtain TgY4 (Fig. 2a). We generated a transgenic line and tested for expression by in situ hybridization on testis sections confirming appropriate site- and stage-specific expression, and by qPCR (Fig. 2 b,c). The breeding test showed 88% female progeny, an unprecedented level of sex ratio distortion with *Smok-Tcr* and a 35% improvement over TgY3 (Fig. 2d, Table 2).

**Figure 2.**
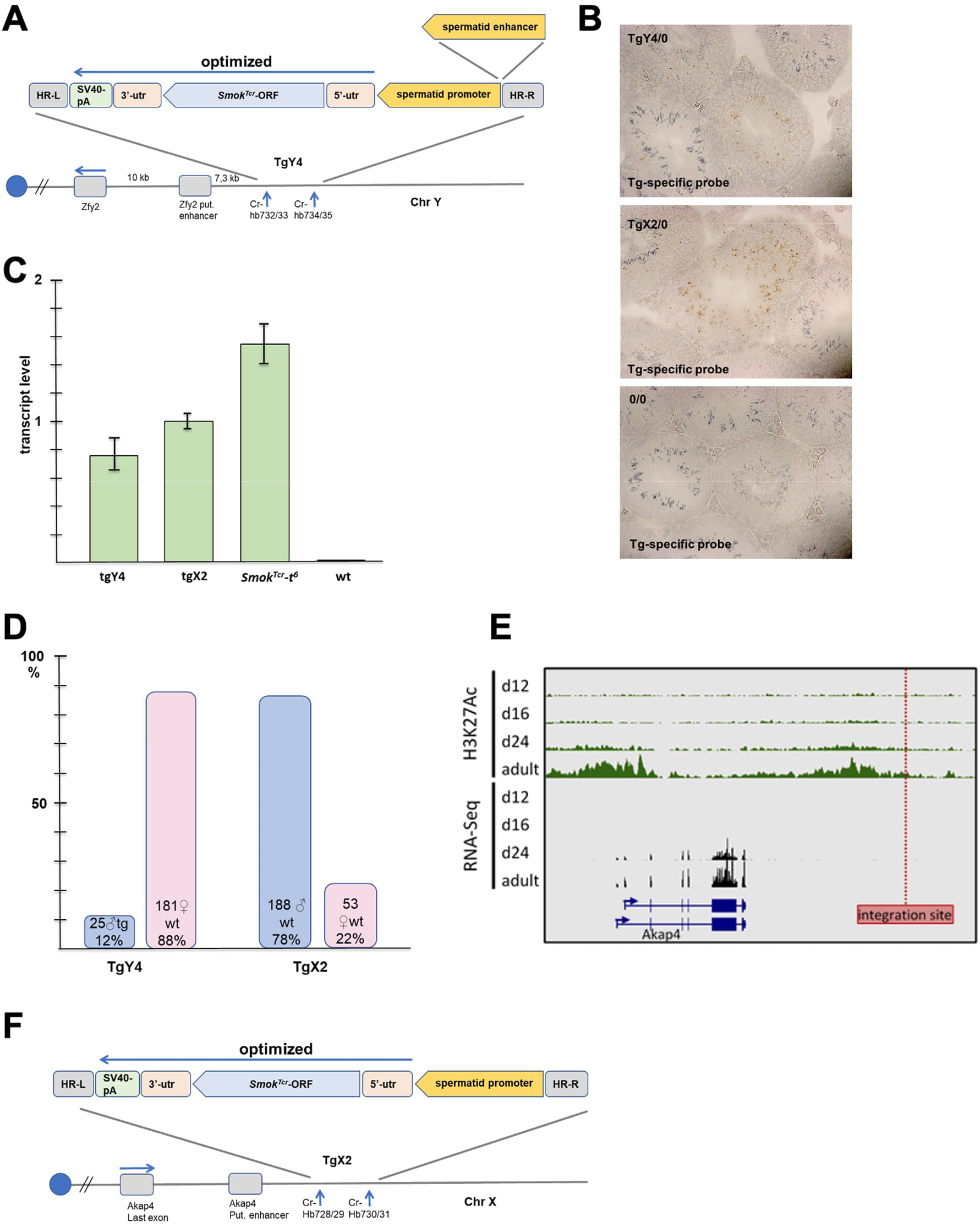
Enhanced sex-ratio distortion by an improved transgene integrated on the Y or the X chromosome. (**A**) Structure of the Y-chromosomal *Smok-Tcr*-transgene tgY4. The spermatid-promoter and enhancer were utilized to drive the expression of an improved Tg cassette integrated in LS-*Zfy2*. (**B**) TgY4 or TgX2 expression detected by in situ hybridization of testis sections is observed in spermatids. (**C**) TgY4 expression detected by q-RT-PCR of testis RNA, in comparison to endogenous *Smok-Tcr*. (**D**) Tg inheritance and sex in offspring from TgY4 or TgX2 males indicate highly efficient sex ratio distortion. (**E**) Identification of a transgene integration site in the vicinity of the *Akap4* locus (LS-*Akap4*). (**F**) Structure of the transgene TgX2 integrated near *Akap4*.

For certain applications, an increased proportion of male offspring is desirable. Therefore, to adapt this approach accordingly we developed an X-integrated, sequence-optimized *Smok-Tcr* transgene, TgX2. Leveraging our RNA-seq and ChIP-seq datasets, we identified a suitable transgene landing site on the X chromosome downstream of *Akap4* (LS-*Akap4*), a region activated late during spermiogenesis (Fig. 2e, Table 1). As a putative enhancer region marked by H3K27Ac levels increasing during spermiogenesis was present near the integration site, we did not include any additional enhancer sequence in our transgene (Fig. 2f).

After generating the transgenic line TgX2, we confirmed appropriate transgene expression (Fig. 2c). Testing for sex ratio distortion demonstrated that TgX2 produced 78% male, non-transgenic offspring (Fig. 2d, Table 2).

### Transgenic males are healthy and fertile

Besides a strong and stable SRD effect the value of a transgenic line highly depends on the health and fertility of the transgenic males. A transgene might affect the wellbeing of a carrier animal by unintended consequences of integration, such as on-or off-target effects during Crispr/CAS mediated integration or, less likely, genomic lesions caused by integration. In addition, transgene expression can be detrimental to the animal’s health. To address issues of general wellbeing, we scored G0 founder animals as well as N1 and N2 backcross animals according to the guidelines of the German Federal Institute for Risk Assessment (BfR). None of our transgenic lines showed a correlation between transgenic status and health issues of animals. Thus, neither our integration procedure, nor the transgenes had a negative impact that manifests in this scoring.

For the applicability of our approach in animal breeding, fertility parameters of transgenic males are of particular interest. Manipulating sperm performance may lead to semisterility, as observed in a recent study (Yosef, Mahata et al. 2023). To determine male fertility we compared the average number of embryos produced in test matings by transgenic males from our SRD lines with those from transgenic lines that did not exhibit SRD (Table 3). No significant differences in embryo numbers were observed between the two groups. Notably, lines expressing optimized *Smok-Tcr*, which exhibited the highest distortion effects, also maintained high fertility (Table 3). These findings indicate that our transgenic males are healthy, fertile, and consistently maintain the SRD effect across subsequent generations.

**Table 3.**
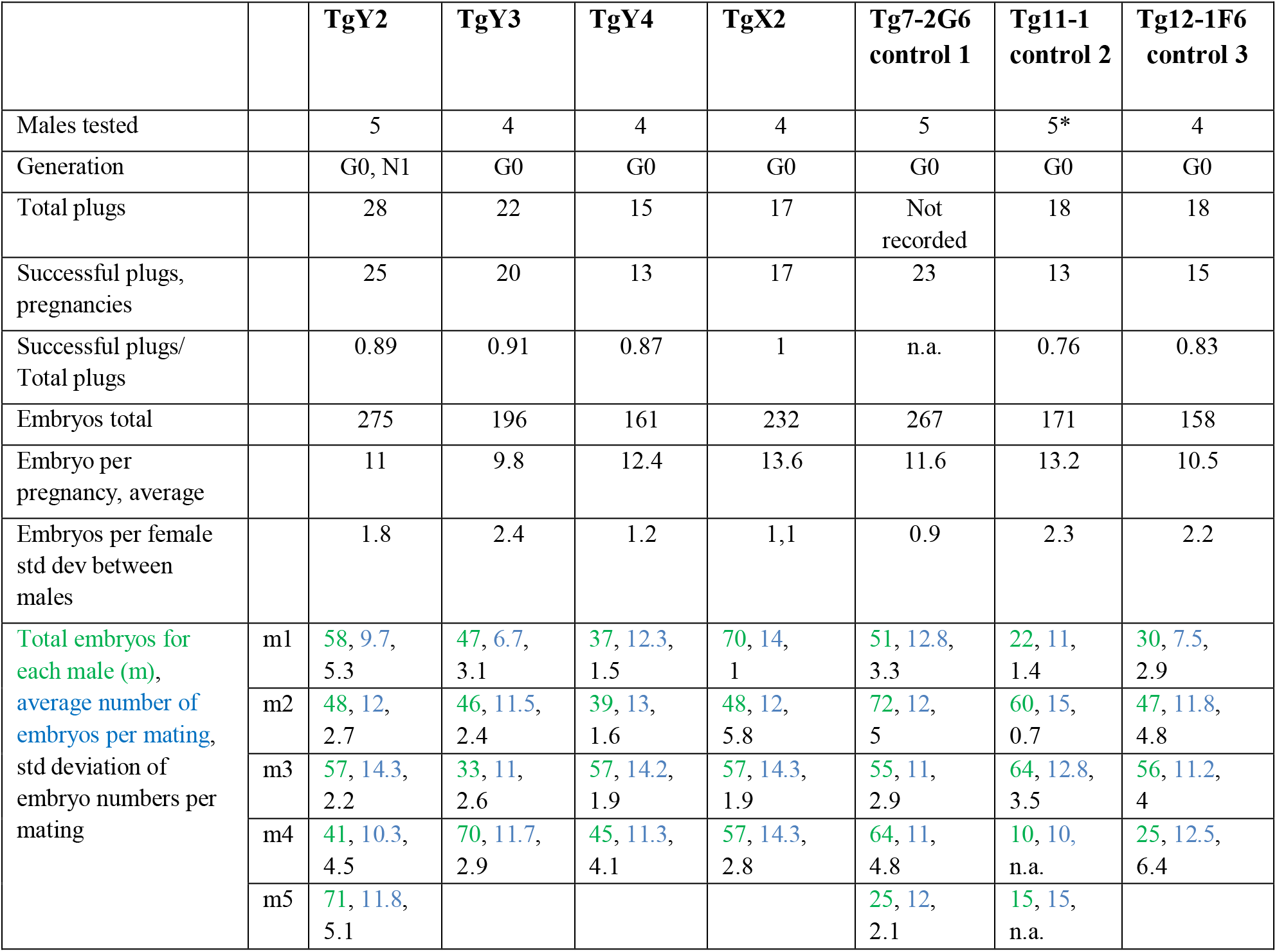
Fertility parameters of transgenic lines showing sex ratio distortion are compared to lines without effect. To test for sex ratio distortion and fertility, each male was mated with a single female for 3 to 5 days per week and plugs were recorded. Females were checked for pregnancy visually and/or by ultrasound examination. Pregnant females were euthanized until d13.5dpc, embryos were dissected and genotyped. Litters for line establishment and maintenance are not included here (note difference in numbers to Table 2); *, Tg11-1: 2 of 5 males (m4, m5) with only one litter.

|  |  | **TgY2** | **TgY3** | **TgY4** | **TgX2** | **Tg7-2G6**<br>**control 1** | **Tg11-1**<br>**control 2** | **Tg12-1F6**<br>**control 3** |
| --- | --- | --- | --- | --- | --- | --- | --- | --- |
| Males tested |  | 5 | 4 | 4 | 4 | 5 | 5* | 4 |
| Generation |  | G0, N1 | G0 | G0 | G0 | G0 | G0 | G0 |
| Total plugs |  | 28 | 22 | 15 | 17 | Not<br>recorded | 18 | 18 |
| Successful plugs,<br>pregnancies |  | 25 | 20 | 13 | 17 | 23 | 13 | 15 |
| Successful plugs/<br>Total plugs |  | 0.89 | 0.91 | 0.87 | 1 | n.a. | 0.76 | 0.83 |
| Embryos total |  | 275 | 196 | 161 | 232 | 267 | 171 | 158 |
| Embryo per<br>pregnancy, average |  | 11 | 9.8 | 12.4 | 13.6 | 11.6 | 13.2 | 10.5 |
| Embryos per female<br>std dev between<br>males |  | 1.8 | 2.4 | 1.2 | 1,1 | 0.9 | 2.3 | 2.2 |
| Total embryos for<br>each male (m),<br>average number of<br>embryos per mating,<br>std deviation of<br>embryo numbers per<br>mating | m1 | 58, 9.7,<br>5.3 | 47, 6.7,<br>3.1 | 37, 12.3,<br>1.5 | 70, 14,<br>1 | 51, 12.8,<br>3.3 | 22, 11,<br>1.4 | 30, 7.5,<br>2.9 |
|  | m2 | 48, 12,<br>2.7 | 46, 11.5,<br>2.4 | 39, 13,<br>1.6 | 48, 12,<br>5.8 | 72, 12,<br>5 | 60, 15,<br>0.7 | 47, 11.8,<br>4.8 |
|  | m3 | 57, 14.3,<br>2.2 | 33, 11,<br>2.6 | 57, 14.2,<br>1.9 | 57, 14.3,<br>1.9 | 55, 11,<br>2.9 | 64, 12.8,<br>3.5 | 56, 11.2,<br>4 |
|  | m4 | 41, 10.3,<br>4.5 | 70, 11.7,<br>2.9 | 45, 11.3,<br>4.1 | 57, 14.3,<br>2.8 | 64, 11,<br>4.8 | 10, 10,<br>n.a. | 25, 12.5,<br>6.4 |
|  | m5 | 71, 11.8,<br>5.1 |  |  |  | 25, 12,<br>2.1 | 15, 15,<br>n.a. |  |

## Discussion

Here we present a genetic system for efficient sex ratio distortion in mice, which can be transferred to farm animals. This is the first time that sex ratio distortion in favor of non-transgenic male offspring has been achieved in a mammal by a single transgene. We selected a suitable promoter/enhancer combination, as well as appropriate landing sites for our transgenes based on histone ChIP-seq experiments in combination with gene expression data. Upon site-specific integration by homologous recombination on the X or Y chromosome, we achieve a sufficiently high level of haploid-specific transgene expression to obtain either predominantly male or female offspring. Corresponding landing sites on the X or Y chromosome of farm animals as well as suitable regulatory elements can be identified with the same criteria, allowing the generation of transgenic farm animals. We have noticed that the transcript level measured for the single copy *Smok-Tcr* gene on chromosome 17 in adult testis is about twice as high as for the transgene construct integrated on the Y chromosome, though the expression level of the endogenous spermatid-specific gene is about ten-fold the level of *Smok2b*, the wildtype ortholog of *Smok-Tcr* (Table 1). This may be due to unfavorable post-transcriptional regulation or could indicate some degree of attenuation of the promoter/enhancer, possibly due to lack of regulatory elements required for full activation in its ectopic environment. However, even though transgene activity appears somewhat compromised, transmission of our less-effective TgY2 and TgY3 is matched only by a single previous *Smok-Tcr* transgene (Tg5-25) randomly (and presumably multi-copy) integrated on an autosome (Herrmann, Koschorz et al. 1999). Our optimized versions TgY4 and TgX2 show superior TRD effects.

Among other strategies developed for sex ratio distortion in mouse, two share features with our approach (Douglas, Maciulyte et al. 2021) (Yosef, Mahata et al. 2023) The first is a *Crispr*- based synthetic lethal system, yielding single sex litters of either males or females, which has been analyzed in great detail on the functional level and with extensive breeding data (Douglas, Maciulyte et al. 2021). The second approach employs *Crispr*i-mediated repression of a gene essential for sperm development (Yosef, Mahata et al. 2023). So far, information on the transgene strategy and its effects provided in this pre-print are sparse and the offspring numbers suggesting high sex ratio distortion are very low. Both approaches have limitations for a practical application. Most importantly, they reduce the fertility of the animal stock. In the synthetic lethal system, half the embryos die resulting in significant reductions of litter size (by 28-39 % due to some compensation). Our system, by keeping the total offspring at a high rate, reduces the proportion of undesired offspring by 38% to 12%. Complete elimination of the less preferred sex (to 0%) achieved by the synthetic lethal system may be desirable for some purposes, whereas in many applications a small proportion of transgenic male progeny required for breeding is preferred. Another disadvantage of the synthetic lethal system is that two transgenic lines have to be maintained separately for animal production. “Escapers” in this system will likely be resistant against the Crispr mediated lethality and thus are not of any use for further breeding. As a side aspect, fully penetrant sex ratio distortion has been regarded as problematic for some applications (Laviola and Adriani 2022).

*t*-haplotype encoded distorter factors can lead to subfertility or sterility of males (Lyon 1986). Although this negative effect is not associated with the responder (Tcr), a possible influence of a transgene acting in the Smok fertility pathway on the breeding performance of males had to be excluded. We therefore analyzed the fertility of the transgenic lines. Generally, matings of laboratory mouse strains show a significant variability in plug rates, proportion of successful plugs (total plugs/pregnancies) and number of embryos and offspring. Not only the performance of a specific breeder male but also other factors are important, such as the age and physical status of the female and conditions in the animal facility. Thus, to mitigate extremes, we based our assessment of fertility on a broader dataset of controls recording data from various (three) other transgenic lines without sex ratio distortion effects, showing low expression of the transgene and tested during different time periods.

In the *Crispr*i system transgenic founder males have growth defects and their fertility is severely compromised (Yosef, Mahata et al. 2023). The IVF procedure necessary to produce sex biased progeny with this system yielded normal litter sizes. However, it is doubtful if the system allows artificial insemination which, together with natural mating is a standard reproductive procedure in farm animals and requires males of good health and semen of good quality. Reduced fertility and growth defects may be caused by the *Crispr*i system’s on- and off target silencing effects, possibly heritable and accumulating. Such considerations are of no concern in our system, restricting the transgene activity to sperm only and impacting on sperm physiology rather than genome and chromatin integrity.

Achieving sex ratio distortion is most desirable in cattle. A variety of specialized breeds for meat production have been developed. Since males gain a lot more muscle in one season than females the former are preferred for beef production, while the latter are commercially less attractive and mostly used for breeding. Our transgene construct inserted on the X chromosome of transgenic bulls would cause sex ratio distortion in favor of non-transgenic males, and thus significantly improve the efficiency in beef production, at the same time avoid the slaughtering of millions of unwanted calves and reduce the CO_2_ burden. Likewise, dairy farming could be improved by avoiding the production of unwanted male calves. Here, the integration of our transgene construct on the Y chromosome would allow generating transgenic bulls as semen producers for artificial insemination of milk cows. Similarly, female offspring is preferred in most farm animals, such as the pig, sheep or goat, and few males are needed for the production of offspring. Therefore, for the latter farm animals a low percentage of Y-transgenic males would suffice for maintenance of a herd/flock.

In summary, our method shows a unique combination of advantages with respect to alternative approaches. In particular when breeding livestock, absence of the transgene in offspring of the desired sex is of great advantage. In several countries such as Australia, New Zealand and the United States, such offspring, although derived from a transgenic parent, is termed “null segregant” and regarded as not genetically modified. Adding to the acceptability of our approach, breeder animals and their offspring are scored healthy. Importantly, transgene activity is maintained over generations and in different genetic backgrounds underscoring the robustness of our system. The fully preserved fertility and fecundity is a particular advantage not present in any of the other approaches published to date. It also suggests that there is scope for further increasing the strength of the distortion effect or tuning it according to particular requirements. Possible strategies include the use of stronger promoters or further optimizations of mRNA stability, translation efficiency or biochemical activity of the protein.

## Materials and Methods

### RNA-Seq

For RNA-isolation, a testis piece, freshly isolated or stored at -80°C was homogenized using a tissueLyser (Qiagen cat. 85220) in a 2 ml Eppendorf tube, 1 min, frequency 30.0.

Total RNA prepared with Trizol (Invitrogen, Thermo Scientific, see below) was DNAse digested and further purified using the RNeasy Micro kit (Qiagen). Any residual genomic DNA was digested on column according to manufacturer’s instructions, with the addition of an extra 1µl of RNase-free DNase I (Roche). The RNA was eluted using RNase-free water, quantified using the Qubit RNA HS Assay and the integrity was verified using Bioanalyzer RNA Pico chips.

Approximately 150-200 ng of total RNA was used for the generation of strand-specific RNA-seq libraries using the ScriptSeq v2 (Epicentre) low input libarary preparation kit according to manufacturer’s instructions. The RNA-seq libraries were quantified using the Qubit high sensitivity DNA assay (Life Technologies) and the size distribution was verified using the DNA HS Bioanalyzer chips (Agilent). Libraries were paired-end sequenced either on a HiSeq 2000 (Illumina) with 2x50bp read length.

### ChIP-seq

We isolated single cells from mouse testes according to (Getun, Torres et al. 2011). Crosslinking of approximately 1x10^6^ cells was performed in PBS essentially as previously described (Koch, Scholze et al. 2017) Cells were washed twice with cold PBS containing 0.05% Triton X-100, pelleted, snap frozen and stored at -80°C until sonication.

Cells were processed using the iDeal ChIP-Seq kit (Diagenode) according to manufacturer’s instructions. Sonification was performed on a Bioruptor Pico (Diagenode) using 3 runs of 10 cycles (30s on, 30s off) in a 4°C water bath. Sheared chromatin was purified and the size distribution was verified using a DNA HS Bioanalyzer chip (Agilent). Approximately 200,000 cells were used for ChIP with the anti-H3K27ac (ab4729, Abcam) antibody.

Approximately 1-5 ng of ChIP DNA was used to generate libraries using the TrueSeq ChIP-Seq kit (Ilumina) with minor modifications. After adapter ligation, a 0.95x volume of AMPure XP beads (Beckman Coulter) was used for a single round of. After the addition of 1 µl primer mix (25 mM each, Primer 1: 5’-AATGATACGGCGACCACCGAG-3’; Primer2: 5’- CAAGCAGAAGACGGCATACGAG-3’) and 15 µl 2x Kapa HiFi HotStart Ready Mix (Kapa Biosystems), amplification was performed for 45 s at 98°C, 5 cycles of [15 seconds at 98°C, 30 s at 63°C and 30 s at 72°C] and a final 1 min incubation at 72°C. The PCR products were purified using a 0.95x volume of AMPure XP beads. The libraries were amplified for further 13 cycles, purified using a 0.95x volume of AMPure XP beads, quantified using the Qubit DNA HS assay and the size was validated using DNA HS Bioanalyzer chips (Agilent). Libraries were sequenced on a NextSeq500 (Illumina) with 1x75bp read length.

### Genome assemblies

Datasets were mapped to the Mus musculus GRCm38/mm10 genome assembly containing chromosomes 1-19, X, Y and M and the refSeq annotations (UCSC) in refflat gtf format.

### Bioinformatic analysis

RNA-seq reads were mapped with TopHat2 (version 2.1.0; (Kim, Pertea et al. 2013) using bowtie (version 1.1.2; (Langmead, Trapnell et al. 2009), providing refSeq annotations and the options ‘–no-coverage-search –no-mixed –no-discordant -g1 –library-type fr-secondstrand’. For visualization, wiggle tracks were generated with BEDTools (version 2.23.0) (Quinlan and Hall, 2010), converted into bigwig format and loaded into the Integrated Genome Browser (Freese, Norris et al. 2016). FPKM’s were calculated using Cuffdiff, part of Cufflinks (version 2.2.1) (Trapnell, Hendrickson et al. 2012) (Trapnell, Williams et al. 2010), with the options ‘-u –no-effective-length-correction -b’.

ChIP-seq data was mapped using bowtie (version 1.1.2) with the options ‘-m 1 -S -y’. We then used MACS (Zhang, Liu et al. 2008) to determine the average fragment length of the sequenced sampled and a custom perl script to elongate the mapped reads to this length. Duplicates were then removed and .wig files were generated using BEDtools (version 2.23.0) (Quinlan and Hall, 2010). The files were converted into bigwig format and loaded into the Integrated Genome Browser (Freese, Norris et al. 2016).

### Identification of spermiogenesis-specific promoters on chromosome X

Genes with a haploid stage-specific expression pattern were selected based on the generated FPKM (Fragments Per Kilobase per Million mapped fragments) values of the staged testis RNA-Seq data. We then isolated genes specific for d16 after birth (p.p.) with an FPKM value of < 2 at d12 and >= 2 at d16 as well as those for d24 with an FPKM value of < 2 at d12 and d16 and >=2 at day 24. We set a cutoff at expression value FPKM < 30.

For the analysis of tissue-specific expression, we obtained CAGE-Seq data from 35 adult mouse tissues (accession E-MTAB-3579; https://www.ebi.ac.uk/biostudies/arrayexpress/studies/E-MTAB-3579) and selected genes with detectable expression in either testis and/or epididymis and removed those with a combined TPM (tags per million) score across all other 33 tissues of less than 1.

To obtain the final list of genomic regions, we then selected the overlap between haploid stage-specific and tissue-specific genes, extracted their respective promoters (-2kb to TSS) and removed those promoters of genes located on autosomes.

### Transgenes

To obtain TgY2 we cloned the promoter fragment of a post-meiotically active gene upstream of the 5’-utr and coding sequence of Tcr-t6 (Herrmann, Koschorz et al. 1999). We attached the 3’-utr of *Smok-Tcr* and a SV40 poly-A signal and flanked the construct with homology regions (HRs) for integration in the vicinity of the *Zfy2* gene. TgY3 is identical to TgY2 but carries in addition a putative enhancer sequence identified by ChIP-seq analyses (see above). We PCR-amplified the putative enhancer sequence by PCR and inserted it 5’-upstream of the promoter. To derive TgY4 we sequence-optimized TgY3. For TgX2 we replaced the *Zfy2* HRs in TgY4 for sequence close to *Akap4*.

### Embryonic stem cell (ESC) culture, genetic engineering and genomic analysis

We carried out ESC culture of G4-F1 hybrid ES cells (gift of A. Nagy)(George, Gertsenstein et al. 2007) on mitotically inactivated embryonic fibroblasts according to standard procedures (Ramirez-Solis, Davis et al. 1993).

We integrated transgenic constructs via homologous recombination, stimulated by Crispr/CAS mediated DNA cleavage. Guide RNA sequences were designed using CRISPOR (http://crispor.tefor.net/) (Concordet and Haeussler 2018). Oligonucleotides were annealed and ligated into the pX330 vector digested with BpiI. (pX330-U6-Chimeric_BB-CBh-hSpCas9 plasmid (Cong, Ran et al. 2013), Gift from Feng Zhang (Addgene plasmid #42230; http://n2t.net/addgene:42230 ; RRID:Addgene_42230).

We confirmed correct, single copy integration of the transgenes by Southern blotting of ES-cell clones using external genomic-or transgene specific probes respectively. Alternatively, we verified single copy integration in breedings. After expansion of Southern-blot positive clones we performed full-length PCR amplification of the integrated transgene using Prime STAR GXL DNA polymerase (TAKARA) with primers outside the integration site and sequenced the PCR product of the integrated transgene.

### Generation, husbandry and transmission test of mouse lines

We expanded correctly targeted ESC clones from frozen 96-well plates to 3.5 cm dishes, froze stocks and, after confirming the sequence of the transgene insertion by PCR (see above), used the clone for ESC aggregation with diploid morulae in the transgenic facility of the MPIMG (Artus and Hadjantonakis 2011). Mouse lines were established by backcrossing to wild type strains such as C57BL/6J.

All animal procedures were in accordance with institutional, state, and government regulations (LAGeSo Berlin, animal licenses G0243/18 and G0098/23 for aggregation experiments and G0309/18 and G0186/23 for G0 animals and the resulting mouse lines).

To determine sex ratio and transgene transmission ratio from the transgenic lines, transgenic males were mated with wild-type females. In the mornings of the following days, females were checked for copulatory plugs and, if plug-positive and pregnant, were sacrificed at 13.5 days after conception. Biopsies of embryos were lysed in Laird’s buffer (Laird, Zijderveld et al. 1991) and genotyped by PCR using transgene-specific primers.

We performed statistical tests of the observed transgene transmission rate in offspring, relative to the expected Mendelian transmission rate of 50%.

### Transgene expression test

Before establishing and breeding the line, a G0 chimeric male was used for transgene expression test. We euthanized sexually mature males by CO_2_ asphyxiation (GasDocUnit, Medres) and isolated testes for RNA isolation and in situ hybridization on paraffin sections using the RNAscope 2.0 HD Detection Kit (Brown) ACD, Bio-Techne) as suggested by the manufacturer except for a longer incubation step in Amp5 solution (60 min instead of 30 min). Staining times were up to 2 hours and adjusted according to signal strength.

We isolated total RNA prepared from testis tissue samples using Trizol (Invitrogen, Thermo Scientific). We removed genomic DNA contamination with the DNA-free kit AM 1906 (Ambion, Invitrogen, Thermo Fisher Scientific). RNA was quantified on a Nano-drop-device (NanoPhotometer, IMPLEN, Version 1.0). We analyzed RNA quality by gel electrophoresis or with Bioanalyzer 2100 (Agilent).

We performed cDNA synthesis for RT-qPCR analysis using 1 μg of testis RNA with the SuperScript reverse transcription system (Invitrogen) or M-MLV Reverse Transcriptase (Promega).

q-PCR Primers were designed with Primer3plus. To generate specific, discriminating primers we used PrimerBLAST and carried out sequence alignment with SnapGene.

## Acknowledgements

The authors thank members of the transgenic facility of the MPIMG, Dr. Ludger Hartmann for supervision of the animal facility as well as Carolin Willke and Sonja Banko for expert animal caretaking.

## Author contribution

Conceptualization, B.G.H.; Construct design, B.G.H., H.B.; Investigation, H.B., F.K., B.L., J.W., G.B., M.S-W., S.W., L.W.; Bioinformatic analysis, F.K.; Writing, H.B., B.G.H., Supervision, B.G.H., H.B.; Funding acquisition, B.G.H.

## Competing interest

BGH, HB and FK have submitted a patent application on the described technology to the European Patent Office.

## Notes

### Funding

- Max Planck Society

